# A novel *C. elegans* respirometry assay using low-cost optical oxygen sensors

**DOI:** 10.1101/2025.05.16.654527

**Authors:** N Dennis, C Gourlay, M Ezcurra

## Abstract

Measurement of the oxygen consumption rate (OCR), or respirometry, is a powerful and comprehensive method for assessing mitochondrial function both *in vitro* and *in vivo*. Respirometry at the whole-organism level has been repeatedly performed in the model organism *Caenorhabditis elegans*, typically using high throughput microplate-based systems over traditional Clark-type respirometers. However, these systems are highly specialised, costly to purchase and operate, and inaccessible to many researchers. Here, we develop a respirometry assay using low-cost commercially available optical oxygen sensors (PreSens OxoPlates®) and fluorescence plate readers (the BMG FLUOstar), as an alternative to more costly standard respirometry systems. This assay uses standard BMG FLUOstar protocols and a set of custom scripts to perform repeated measurements of the *C. elegans* OCR, with the optional use of respiratory inhibitors or other interventions. We validate this assay by demonstrating the linearity of basal OCRs in samples with highly variable numbers of animals, and by examining the impact of respiratory inhibitors with previously demonstrated efficacy in *C. elegans*: carbonyl cyanide 4-(trifluoromethoxy) phenylhydrazone (a mitochondrial uncoupler) and sodium azide (a Complex IV inhibitor). Using this assay, we demonstrate that the sequential use of FCCP and sodium azide leads to an increase in the sodium azide-treated (non-mitochondrial) OCR, indicating that the sequential use of respiratory inhibitors, as standard in intact cell respirometry, may produce erroneous estimates of non-mitochondrial respiration in *C. elegans* and thus should be avoided.

## Introduction

The mitochondria are central to cellular health, performing critical functions in metabolism, ion homeostasis, cell signalling, and ATP synthesis, among many other processes (reviewed in references 1,2). Consequently, mitochondrial dysfunction is associated with a diverse spectrum of human diseases, most notably genetic mitochondrial diseases^3^, but also common and increasingly prevalent conditions such as Alzheimer’s disease, obesity, and type 2 diabetes^4–6^.

Mitochondrial health can be examined via numerous methods, including the microscopic evaluation of mitochondrial morphology^7^, estimation of the mitochondrial membrane potential^8^, bioluminescence-or fluorescence-based ATP assays^9–11^, and PCR-based assays of mtDNA copy number^12^. However, while each of these methods provide useful metrics of mitochondrial function, they fail to provide a complete picture of mitochondrial health.

In contrast, measurement of the oxygen consumption rate (OCR), or respirometry, is the most direct and comprehensive method of examining cellular metabolism and mitochondrial function^13^. As the vast majority of oxygen consumption occurs via the mitochondrial electron transport chain (ETC)^14^, combining OCR measurements with specific mitochondrial substrates and ETC inhibitors can provide informative readouts of mitochondrial function, such as the proportion of ATP synthase-linked respiration, the maximal respiration rate, and the level of non-mitochondrial oxygen consumption^15^. OCRs can be measured in isolated mitochondria, intact or permeabilised cells, as well as 3D models (tissues, organoids or whole organisms), providing scope for the assessment of mitochondrial function in multiple contexts^13^.

While respirometry in isolated mitochondria greatly simplifies the interpretation of OCR data, experiments in whole organisms uniquely reflect all factors influencing physiological respiratory activity, preserving the integrity of the mitochondrial network^16^, their interactions with other cellular components, and the influence of intercellular signalling processes^17,18^. This approach to respirometry has been repeatedly performed in the nematode *Caenorhabditis elegans*^19–29^, a model organism which has been used to study mitochondrial dynamics^30^, mitochondrial retrograde signalling^31^ and mitochondrial disease^32^. Respirometry studies of *C. elegans* have elucidated the impact of genetic defects in electron transport chain subunits, fission-fusion dynamics and mitophagy^19^, examined sex- and development-specific respiratory profiles^27,29^, and evaluated OCRs as a sub-lethal endpoint for toxicity screening^28^.

Respirometry is generally performed using one of two categories of instrumentation: large, chamber-based Clark-type electrode systems, such as the Oroboros Oxygraph-2k; and small, specialised microplate-based systems using oxygen-sensitive phosphors, such as Agilent Seahorse extracellular flux (XF) analysers^13^. Respirometry studies in *C. elegans* have been performed using both Clark-type and microplate-based respirometers^19,21,25,26^, but the large requirement for biological material and low throughput associated with Clark-type respirometry^13^ has led to a clear preference for the latter, as reflected in numerous *C. elegans* protocols outlining the use of the 8-well, 24-well and 96-well Seahorse XF analysers^20–24^. While the relatively low requirement for biological material and high throughput of microplate-based systems make them ideally suited for work with *C. elegans*, these systems come with a major drawback in the form of high upfront and ongoing costs^13,33^. For many researchers, these costs are likely prohibitive, rendering microplate-based respirometry inaccessible.

To make *C. elegans* respirometry experiments more accessible, we developed a respirometry assay using low-cost commercially available optical oxygen sensors and fluorescence plate readers. Our assay utilises OxoPlates® (manufactured by PreSens Precision Sensing), 96-well microplates featuring phosphorescent oxygen sensors. These systems have been applied to high throughput investigations of oxygen consumption in bacteria, yeast, and mammalian cell culture, often as a metric of cell viability^34–37^, but have seen limited and variable usage in *C. elegans*^38,39^. Each OxoPlate sensor consists of two dyes embedded in a thin polymer matrix: an indicator dye, whose phosphorescence intensity (I_indicator_) is inversely proportional to the oxygen concentration; and a reference dye (I_reference_), whose phosphorescence intensity is independent of the oxygen concentration. Oxygen concentrations are calculated from internally referenced sensor responses 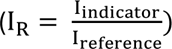, using I_R_ signals derived from oxygen-saturated (k_100_) and oxygen-free (k_0_) calibration solutions, where 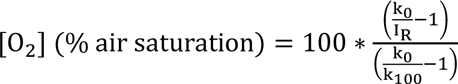. The only hardware requirement associated with these sensors is a plate reader capable of performing fluorescence intensity or, ideally, time-resolved fluorescence measurements, using dual kinetics and bottom optical sensors.

The assay described here was developed for BMG FLUOstar fluorescence plate readers but could likely be adapted for other readers with similar functionality. Our assay utilises the “kinetic windows” program of the BMG FLUOstar to perform sequential measurements of the *C. elegans* OCR in multiple windows of measurement (oxygen consumption) and linear shaking (re-oxygenation). We designed a simple script to loop several of these programs in sequence with the option of pausing the script for the manual addition of respiratory inhibitors or other interventions, and an accompanying data analysis script to automatically calculate OCRs from linear regressions fit to each measurement window (represented graphically in Figure 1).

**Figure 1.**
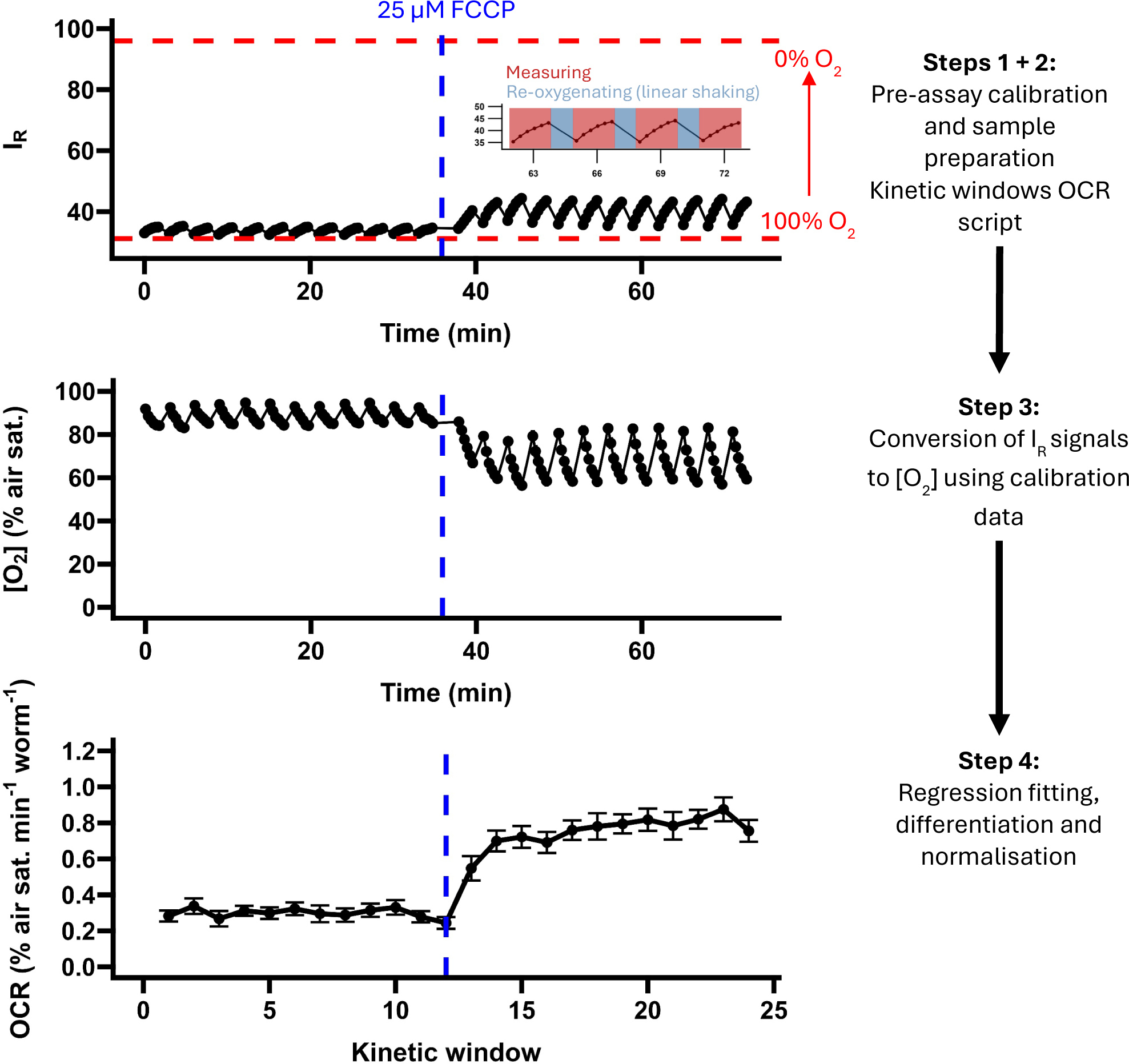
Graphical summary of the assay. **Step 1**, Sample preparation (collection, washing and loading of *C. elegans* samples) and pre-assay sensor calibration using oxygen-saturated (100 % O_2_) and oxygen-free (0% O_2_) calibration solutions. **Step 2**, A kinetic windows measurement protocol is looped by an OCR tracing script, generating internally referenced (I_R_) signal data with characteristic increases over a 100 s measurement period (as oxygen is consumed by the samples) and declines (as a 1-minute 700 RPM linear shaking period re-oxygenates the wells; see inset). **Step 3**, I_R_ signals are converted into oxygen concentrations (as a percentage of air saturation) using the two-point calibration data from step 1 (denoted with red dashed lines). **Step 4**, an OCR analysis script fits linear regressions to each period of oxygen consumption (kinetic window), extracts the regression coefficient, multiplies the result by −1, and normalises the resulting oxygen consumption rate (OCR) using a normalisation parameter (here the number of animals per well). The data presented here are mean I_R_, [O_2_] and OCR (± SE) traces from an experiment using an optimal concentration of the uncoupler carbonyl cyanide 4-(trifluoromethoxy) phenylhydrazone (FCCP; re-produced from Figure 3; N = 19 wells; approximately 20 day 1 adult-stage animals per well).

To validate this assay, we first measured basal OCRs in wells with highly variable numbers of day 1-adult stage animals to establish a range of qualitative accuracy, and proceeded to examine the effects of two-well characterised respiratory inhibitors in a set of titration experiments: carbonyl cyanide 4-(trifluoromethoxy) phenylhydrazone (FCCP; a mitochondrial uncoupler) and sodium azide (a Complex IV inhibitor^40^). We then tested whether these drugs can be used sequentially, as standard in intact cell respirometry^13^, using optimal concentrations derived from our titration experiments. We show that this assay can be used to accurately estimate the *C. elegans* OCR at a similar scale to Seahorse XF24 and XF96 respirometers (3-72 animals per well) and can capture the effects of FCCP and sodium azide with optimal concentrations highly similar to those reported in Seahorse XF analyser-based studies of *C. elegans* (25 µM FCCP, 24 mM sodium azide). We also show that treatment with FCCP leads to an elevation of the sodium azide-treated (non-mitochondrial) OCR, supporting a previous Seahorse XF analyser study^19^ and indicating that accurate measurements of the *C. elegans* non-mitochondrial OCR must be obtained separately from measurements of the maximal OCR.

## Materials and Methods

### *C. elegans* strains and maintenance

Bristol N2 (wild type) *C. elegans* were obtained from the Caenorhabditis Genetics Centre (CGC, University of Minnesota). Animals were maintained on Nematode Growth Media (NGM) plates seeded with a thin lawn of *Escherichia coli* OP50 as previously described^41^. Age-synchronised populations were obtained via sodium hypochlorite-based bleaching protocols followed by overnight incubation in M9 at 20 °C with gentle rocking^42^. Synchronised L1s were transferred onto NGM plates seeded with *E. coli* OP50 and were grown at 20 °C until day 1 of adulthood (72 h later) for all experiments.

### BMG FLUOstar and OxoPlate preparation

All experiments were conducted using a BMG FLUOstar Omega plate reader at 25 °C. OxoPlate sensor readouts were obtained using dual kinetic time-resolved fluorescence protocols set up in accordance with PreSens’ documentation. Indicator-dye fluorescence (I_indicator_) was measured with an excitation wavelength of 544 nm and an emission wavelength of 655 nm; reference-dye fluorescence (I_reference_) was measured with an excitation wavelength of 544 nm and an emission wavelength of 590 nm. Additional protocol parameters are described in Table 1.

**Table 1.** Basic time-resolved fluorescence protocol parameters.

| Multichromatics | Integration time | Settling time | Flashes well<sup>-1</sup> |
| --- | --- | --- | --- |
| 544 ex / 590 em (indicator) | 12 – 512 μs | 0.3 s | 20 |
| 544 ex / 655 em (reference) |  |  |  |

In its default configuration, the BMG FLUOstar cannot perform time-resolved fluorescence measurements using its bottom optical sensors, which is a requirement for accurately reading the OxoPlate sensors. As such, prior to all experiments the light guides were reconfigured as shown in Figure S1, in accordance with PreSens’ instructions.

All experiments were conducted using freshly opened OxoPlates or the unused wells of previously opened OxoPlates that were maintained in accordance with PreSens’ instructions. At least twenty minutes prior to an assay, the OxoPlate sensors were hydrated with a small volume of M9 (80 µL for titration and sequential drug treatment experiments; 20-95 µL for sample size experiments).

### OxoPlate sensor calibration

Each batch of OxoPlates were calibrated using an oxygen-saturated calibration solution (Cal100) and an oxygen-free calibration solution (Cal0). The Cal100 solution was prepared by vigorously shaking Milli-Q water for two minutes in a container with a large air phase. This solution was then rested, uncapped, for 1 minute prior to use. The Cal0 solution was prepared as a 1 % w/v solution of sodium sulfite in a container with a minimal air phase. These solutions were transferred in 100 µL aliquots to the OxoPlate wells in technical quadruplicate. The wells containing Cal0 solution were sealed with 100 µL mineral oil to minimise oxygen ingress.

Calibration measurements were preceded by a 30-minute incubation in the plate reader to ensure the contents of the wells had equilibrated to the internal temperature of the reader. In accordance with PreSens’ documentation, one calibration was used for all subsequent experiments using the same batch of OxoPlates. To verify whether the media, solvents and drugs used in our validation experiments warranted separate calibrations, we performed these calibrations in M9 buffer, with 0.2 % DMSO (FCCP solvent), and with optimal concentrations of respiratory inhibitors derived from our dose-response analyses. However, we noted negligible and non-significant alterations to I_R_ signals in all these conditions (Table S1). As such, all presented oxygen concentrations were calculated using water-based calibrations as described above.

### Sample preparation and loading

*C. elegans* sample preparation followed a modified version of previously published *C. elegans* respirometry protocols^20,21^. Populations of approximately 300-500 age-synchronised day 1-adult stage *C. elegans* were washed from NGM plates into 15 mL falcon tubes with M9, allowed to settle without centrifugation and the supernatant was discarded. The samples were then washed with M9 and placed on a rocker for 10 minutes to allow the animals to clear their gut contents of residual bacteria. Following this incubation, the samples were once again allowed to settle without centrifugation and washed with M9. This was repeated at least twice or until all visible eggs, offspring and bacterial debris had been cleared. After the final wash, the population density of the solution was estimated by calculating the average number of worms in five 20 µL droplets using a dissecting microscope. The solution was then adjusted to an approximate population density of 1 worm µL^-1^ with M9 and transferred to the hydrated OxoPlate wells to a final well volume of 100 µL. Approximately 20 animals (i.e. 20 µL of the worm solution) were transferred into each well for all titration and sequential drug treatment experiments. Sample size experiments used highly variable numbers of animals and adjusted the amount of worm solution accordingly (5-80 µL); for measurements with 80 + animals, the worm solution was concentrated to 2-3 worms µL^-1^ and transferred in 80 µL aliquots.

### Drug preparation and treatments

Carbonyl cyanide 4-(trifluoromethoxy) phenylhydrazone (FCCP) stocks were prepared in dimethyl sulfoxide (DMSO) at 500 X final concentration. Sodium azide stocks were prepared in Milli-Q water at 33 X final concentration.

### OCR measurement protocols

OCRs were measured by tracking oxygen consumption over multiple intervals using the BMG FLUOstar’s kinetic windows program. The basic OCR protocol featured four kinetic windows, with readings taken every 20 seconds for 100 seconds, each preceded by a one-minute period of linear shaking at 700 RPM to re-oxygenate the measurement wells (Table 2). With these protocol parameters, maintaining a cycle time of 20 seconds allowed us to measure a maximum of 10 wells per protocol, and brought the length of each protocol to a total of 12 minutes.

**Table 2.**
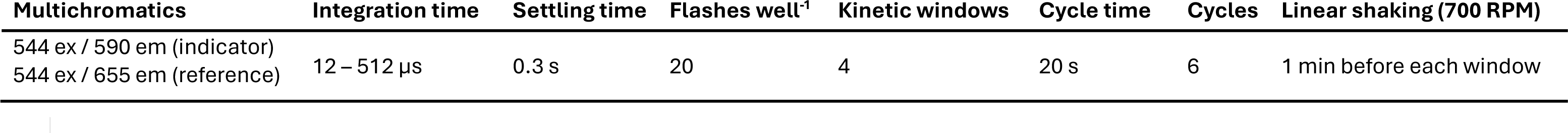
Time-resolved fluorescence protocol parameters for oxygen consumption rate measurements.

| Multichromatics | Integration time | Settling time | Flashes well<sup>-1</sup> | Kinetic windows | Cycle time | Cycles | Linear shaking (700 RPM) |
| --- | --- | --- | --- | --- | --- | --- | --- |
| 544 ex / 590 em (indicator) | 12 – 512 μs | 0.3 s | 20 | 4 | 20 s | 6 | 1 min before each window |
| 544 ex / 655 em (reference) |  |  |  |  |  |  |  |

All assays were performed using a custom BMG FLUOstar script that looped these protocols in sequence, with every third protocol separated by a pause to allow for the manual injection of respiratory inhibitors (for further details see OCR protocol script in the Supplementary Information). For each assay, the number of protocol loops were altered, with sample-size tests assessed with two, drug titrations assessed with six (three control and three drug-treated), and sequential drug treatment experiments assessed with nine (three control and six drug treated). Following each assay, the contents of each well were extracted using pipette tips coated in 0.01 % Triton X-100 in M9, and the number of animals residing in each well were counted by eye using a dissecting microscope.

### Data analysis and statistics

Internally referenced (I_R_) sensor responses were calculated from reference- and indicator-dye phosphorescence using the BMG MARS data analysis software, where 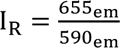 Oxygen concentrations were calculated using calibration constants derived from the average I_R_ signals of oxygen-free (k_0_) and oxygen-saturated (k_100_) calibration solutions (see OxoPlate sensor calibration), where: 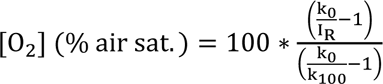.

OCRs were quantified in each kinetic window by fitting linear regressions to the first minute of oxygen consumption, extracting the regression coefficient, and multiplying the result by −1, i.e. assuming [O_2_] ≈ at + b, OCR ≈ −a. OCRs were then normalised to the number of worms within each well. This process was automated using a custom data analysis script written for R v. 4.3.3 (see Automated OCR calculation script in the Supplementary Information).

Parameters of respiratory function were calculated as follows: basal (i.e. routine or untreated) OCRs were calculated as the average OCR prior to treatment with any respiratory inhibitor; maximal OCRs were calculated as the average of the highest three OCRs following treatment with FCCP, with the first three measurement windows excluded; and non-mitochondrial OCRs were calculated as the average OCR following treatment with sodium azide.

All statistics were calculated using R v. 4.3.3. The effects of sample size on per-well OCRs were analysed via linear regression. Unless stated otherwise, all remaining data were analysed using ANOVA with post-hoc Tukey’s Honestly Significant Differences (TukeyHSD) tests. All graphs were produced using the R package *ggplot2*^43^ and all data presented in graphs and tables display the mean ± standard error. All experiments were repeated at least twice, and the data were pooled for analysis.

## Results and Discussion

### Sample size analysis

To validate our assay, we first examined the relationship between basal OCRs and the quantity of worms within each OxoPlate well, as a linear relationship between basal OCRs and the population densities of the sample wells would establish qualitative accuracy over a defined range. Therefore, we measured OCRs in wells containing highly variable numbers of worms (3-280 animals per well). Our results indicated that basal OCRs were linear with population density between 3 and 72 animals per well (R^2^ = 0.92, F_1, 30_ = 334.5, P < 0.0001; Figure 2A), with an average worm-normalised OCR of 0.248 % air sat. min^-1^ worm^-1 (^Figure ^2^B). OCRs sharply declined beyond this range (112-280 animals per well), with an average worm-normalised OCR of 0.01 % air sat. min^-1^ worm^-1^ (F_1, 38_ = 91.2, P < 0.0001 relative to worm-normalised OCRs in the linear range; Figure 2B), as the high rate of oxygen consumption exceeded the capacity of the linear shaking period to sufficiently re-oxygenate the wells (Figure S2A), resulting in OCR-limiting hypoxia (F_1, 38_ = 335.5, P < 0.0001; Figure S2B). These results demonstrate that our assay can be used to estimate the *C. elegans* OCR with relatively small quantities of worms, making sample preparation straightforward. This requirement for biological material (3-72 day 1 adult-stage animals per well) is similar to the amount required for Seahorse XF96 and XFp respirometers (∼ 2-25 day 1 adult-stage animals per well^21^) and Seahorse XF24 respirometers (∼50 day 1 adult-stage animals per well^24^), and is far lower than the requirement for Clark-type respirometry (300-1000 day 1 adult-stage animals per chamber^25,44^).

**Figure 2.**
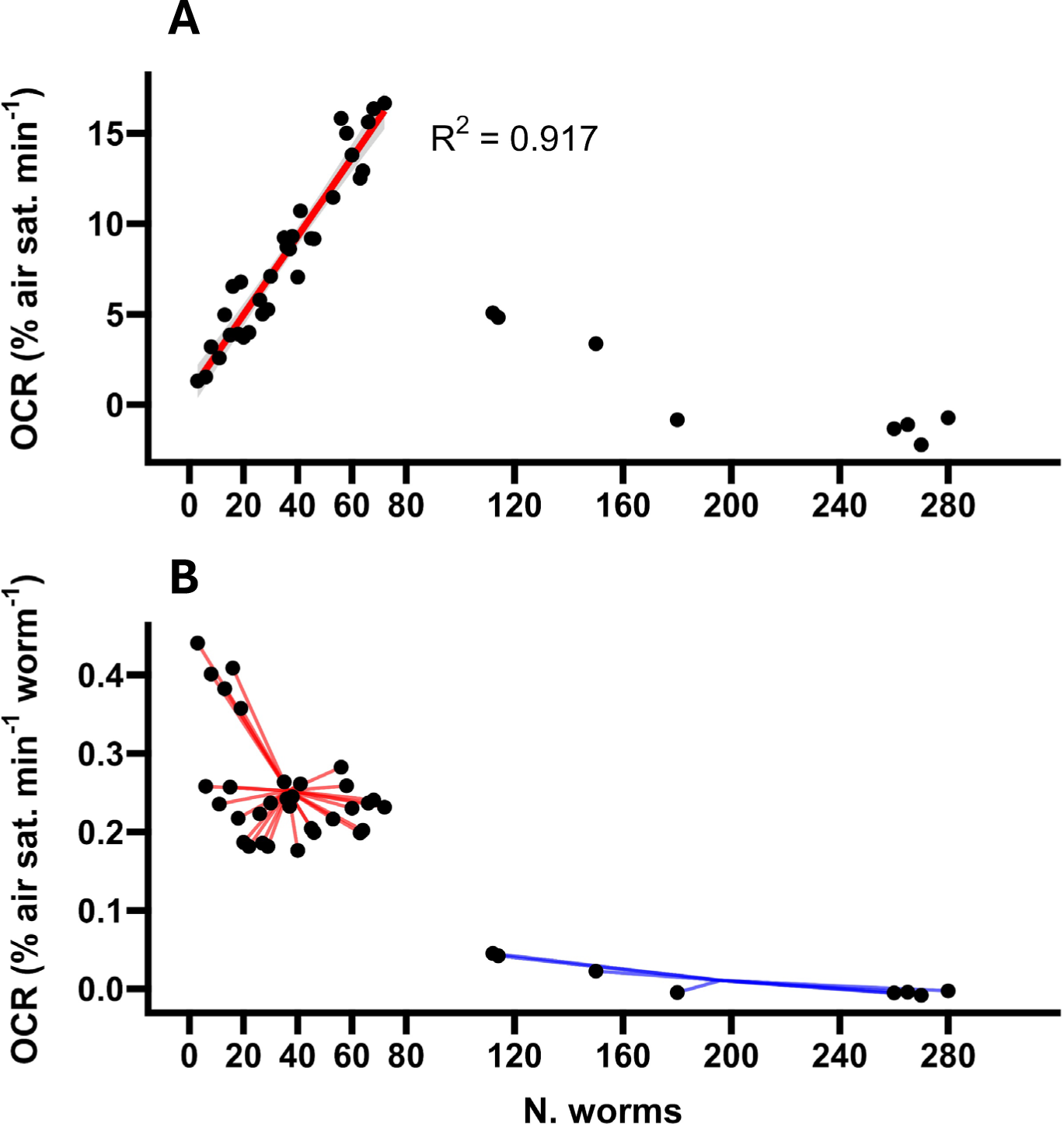
The impact of sample size on basal OCRs. **A,** Average oxygen consumption rates (OCRs) in wells containing N = 3 to N = 280 worms. The linear regression is fitted to OCR data from 3 - 72 worms. **B,** Average worm-normalised OCRs in wells containing N = 3 to N =280 worms. The line segments within each group are drawn from each data point to the group-mean OCR (mean OCR N = 3-72: 0.248 % air sat. min^-1^ worm^-1^; mean OCR N = 112-280: 0.01 % air sat. min^-1^ worm^-1^). N = 40 wells.

### Responses to respiratory inhibitors

Next, we tested whether our assay could capture alterations in respiratory activity caused by respiratory inhibitors with previously demonstrated efficacy in *C. elegans*. Respirometry in intact cells is typically performed with the mitochondrial ATP synthase inhibitor oligomycin to assess ATP synthase-coupled respiration, the protonophore FCCP to assess maximal (uncoupled) respiration, and a combination of the Complex I inhibitor rotenone and Complex III inhibitor antimycin A to assess non-mitochondrial respiration^15^. However, with the exception of FCCP these compounds are ineffective in *C. elegans*, likely due to poor diffusion through the cuticle^19,21^.

Instead, respirometry experiments in *C. elegans* generally use FCCP to assess maximal respiration, and the Complex IV inhibitor sodium azide to assess non-mitochondrial respiration^19–23^. The non-specific ATP synthase inhibitor dicyclohexylcarbodiimide (DCC)^45^ has been used to examine mitochondrial ATP synthase-linked respiration in *C. elegans*^19,20,46^, but its action requires a long incubation period (approximately 60 minutes)^19^, and at least one study has noted a failure to observe any detectable OCR response following treatment with DCC^22^. We therefore focused primarily on examining the effects of FCCP and sodium azide.

To determine the optimal concentrations of these drugs, we performed a set of titration experiments. Concentrations of 25 µM FCCP and 12 mM sodium azide were taken as starting points based on similar estimates of optimal concentrations reported in Luz et al.^20^, and a range of concentrations around these values, in log_2_ scale, were examined, with responses assessed following a period of basal respiration. Based on our previous sample size results, and anticipating the eventual sequential use of FCCP and sodium azide, we opted to use 20 worms per well for all subsequent experiments, reasoning that this would allow OCRs to fluctuate significantly while remaining within the linear OCR range shown in Figure 2A.

As expected, treatment with FCCP significantly altered the maximal respiration rate (F_7, 84_ = 12.1, P < 0.0001; Figure 3A), with a clear peak response observed at 25 µM (Figure 3B). Treatment with 50 µM FCCP did not further alter the maximal OCR relative to the 25 µM treatment (P = 0.945; TukeyHSD), while treatment with 100 µM FCCP induced a significant decline (P < 0.0001; TukeyHSD). As FCCP is water insoluble, it is possible that this was partially caused by the precipitation of FCCP over the course of the assay. However, it is also well-known from studies in intact cells that excessive concentrations of protonophores such as FCCP can lead to a severe drop in respiratory activity, likely resulting from a reduction in substrate import caused by the collapse of the proton-motive force^47^, which may have contributed to the repression of maximal OCRs observed here. Our estimated optimal concentration of 25 µM matches that described in Luz et al.^20^ and is within the range described in *C. elegans* Seahorse respirometry protocols, which commonly use concentrations between 10-25 µM^20–24^. Moreover, the responses we found here are of approximately the same relative magnitude (1.92-fold increase relative to DMSO controls at 25 µM FCCP) to those reported elsewhere (approximately 2-fold increase relative to DMSO controls in Koopman et al.^21^ and Luz et al.^19^), indicating that our assay is able to accurately capture maximal respiration in *C. elegans*.

**Figure 3.**
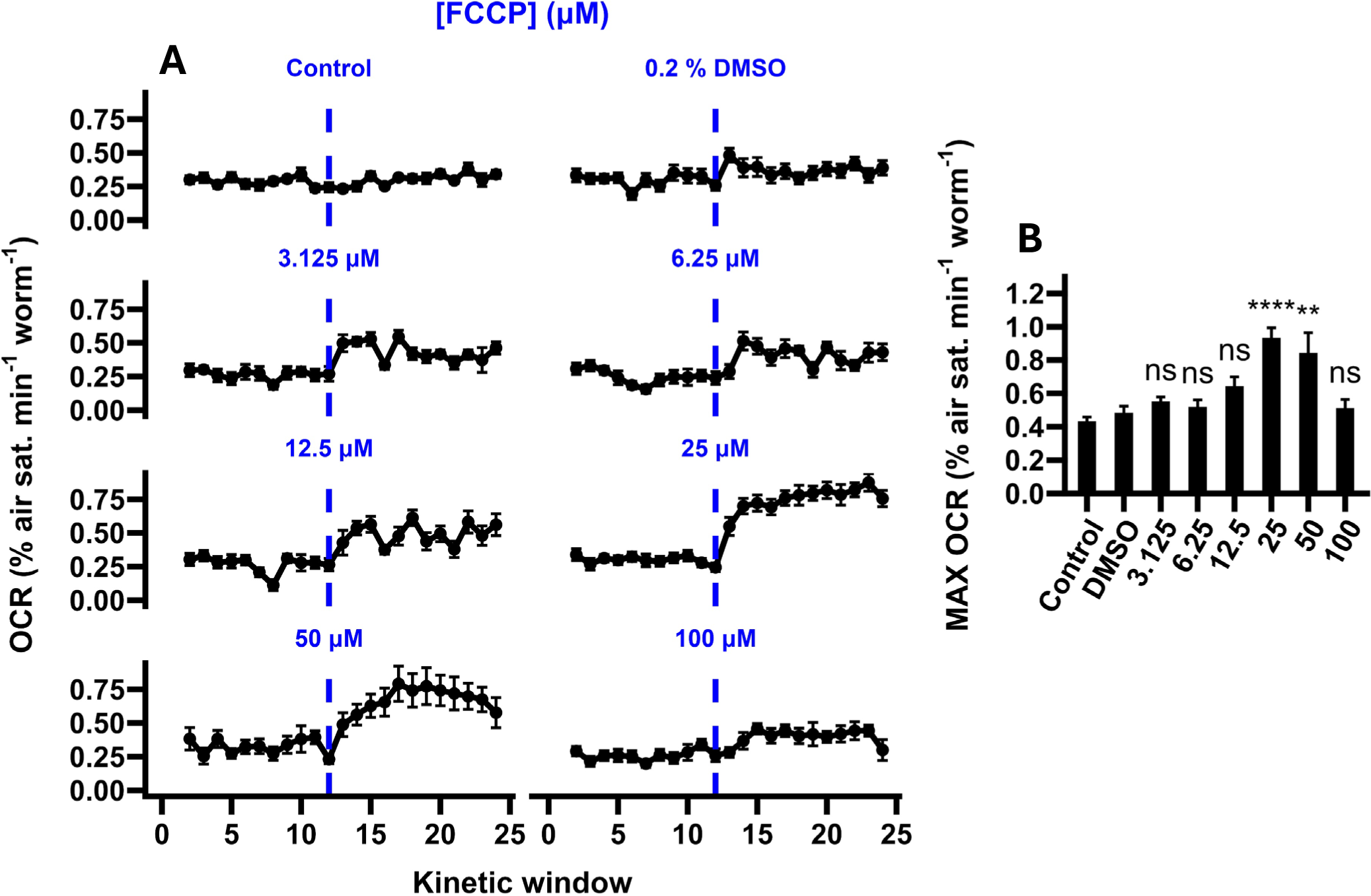
The impact of FCCP on maximal OCRs. **A,** Oxygen consumption rate (OCR) traces of *C. elegans* samples supplemented with varying concentrations of FCCP (dashed lines). Untreated control samples received no drug or solvent treatment, and solvent controls received an injection of DMSO (final concentration 0.2 % v/v). **B,** Maximal OCRs calculated from OCR traces presented in A. Maximal OCRs were calculated as the average of the highest three OCR measurements following FCCP treatment (kinetic windows 16-24). Annotations represent the results of post-hoc TukeyHSD tests relative to DMSO controls: ****, P < 0.0001; **, P < 0.01; ns, not significant (P > 0.05). N = 9-19 wells.

Treatment with sodium azide also strongly altered OCRs (F_8, 44_ = 14.15, P < 0.0001; Figure 4A), with significant reductions observed at concentrations greater than 6 mM (Figure 4B). Relative to the 6 mM treatment, higher concentrations were not associated with further repression of the OCR (all P > 0.938; TukeyHSD), but were associated with a shorter response time (the number of kinetic windows required to reach the average basal-normalised non-mitochondrial OCR; F_3, 20_ = 5.34, P = 0.0072; Figure 5A), with the fastest responses observed in the 24 mM and 48 mM treatments (Figure 5B). As the 48 mM treatment did not further reduce the response time relative to the 24 mM treatment (P = 0.86; TukeyHSD), we concluded that the optimal concentration of sodium azide was 24 mM. This concentration is within the range described in C. elegans Seahorse respirometry protocols, which commonly use concentrations between 10 mM and 50 mM^20–23^.

**Figure 4.**
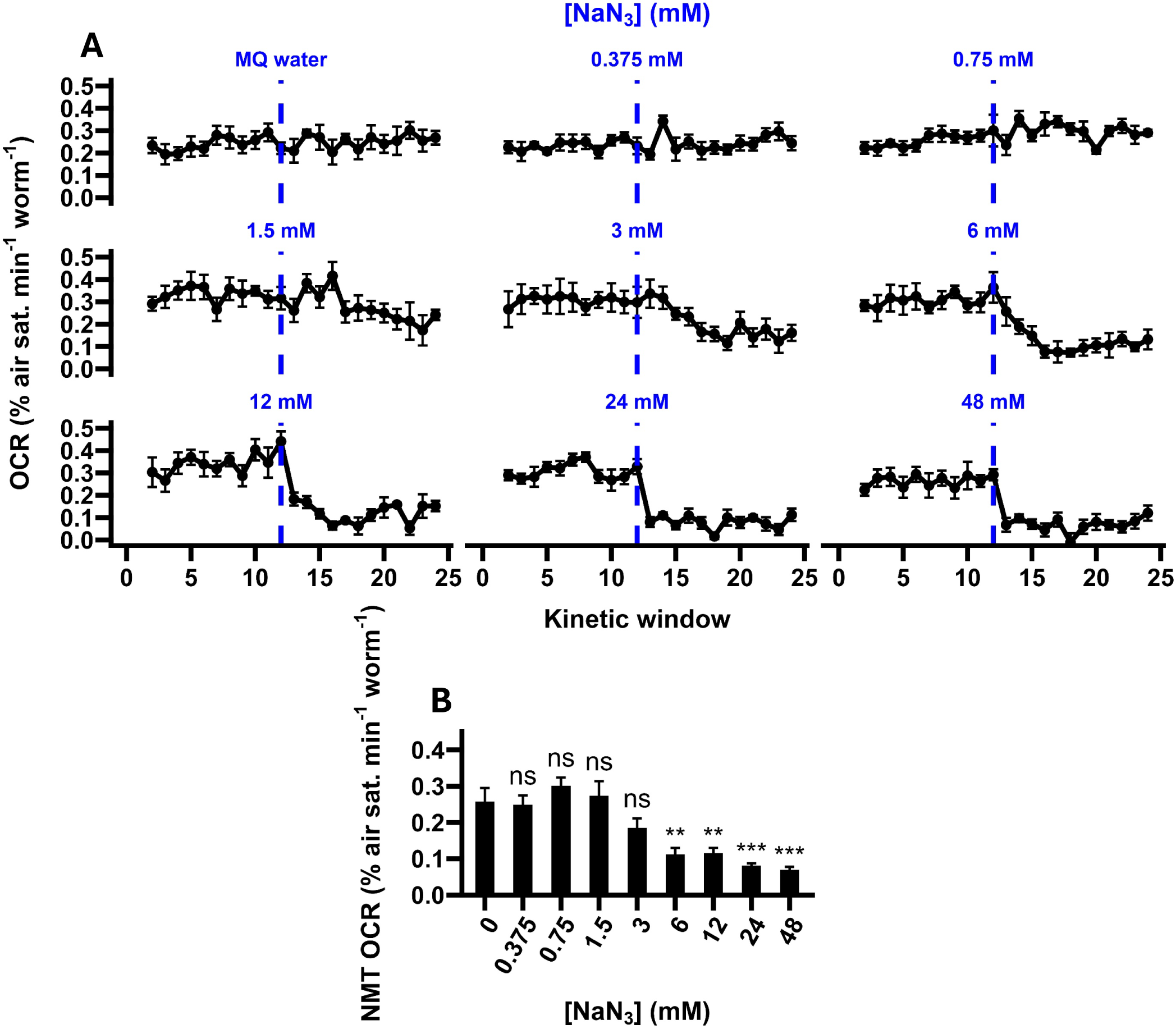
The impact of sodium azide on non-mitochondrial OCRs. **A,** Oxygen consumption rate (OCR) traces of *C. elegans* samples treated with varying concentrations of sodium azide (NaN_3_; dashed lines). Solvent controls received an injection of Milli-Q (MQ) water. **B,** Non-mitochondrial (NMT) OCRs calculated from the OCR traces presented in A. Non-mitochondrial OCRs were calculated as the average OCR following sodium azide treatment (kinetic windows 13-24). Annotations represent the results of post-hoc TukeyHSD tests relative to MQ water controls: ***, P < 0.001; **, P < 0.01; ns, not significant (P > 0.05). N = 5-7 wells.

**Figure 5.**
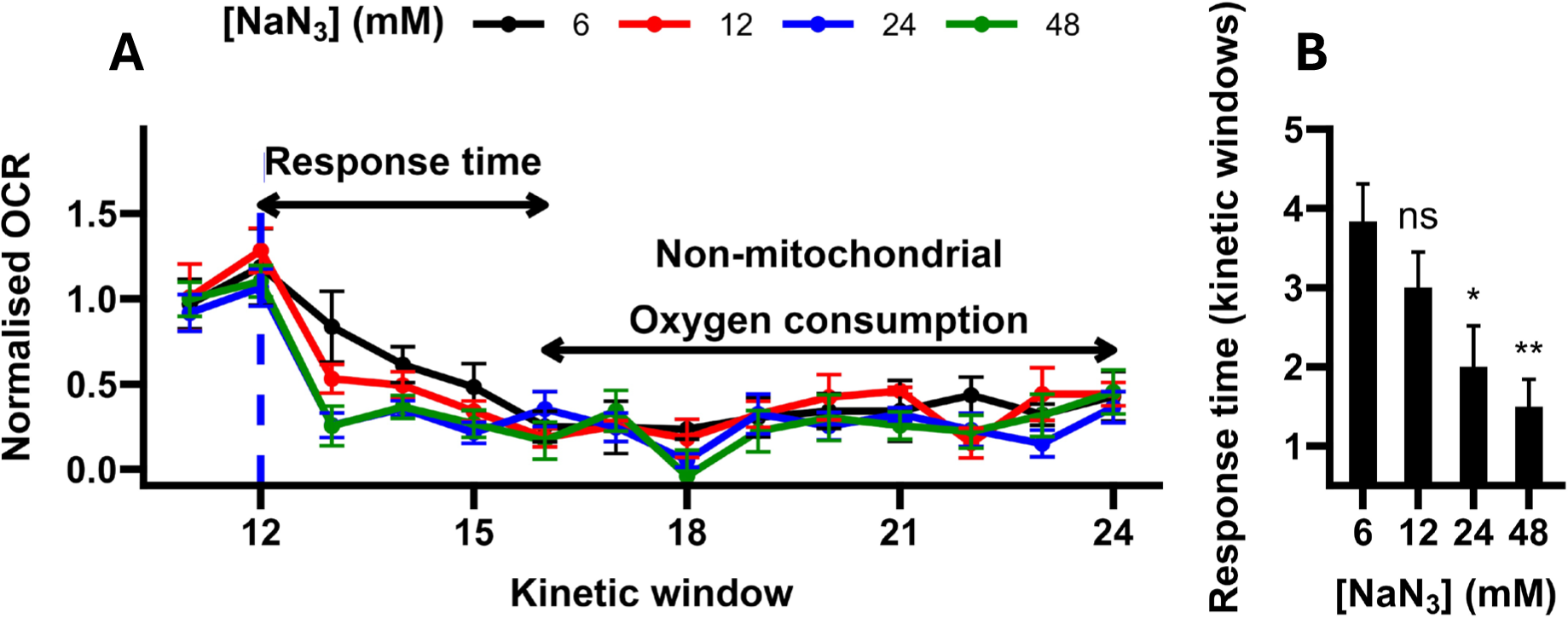
Sodium azide response times. **A,** Basal-normalised oxygen consumption rate (OCR) of samples treated with 6-48 mM sodium azide (NaN_3_; dashed line). OCRs were normalised using the sample-mean basal OCR (kinetic windows 1-12). **B,** Average response times. Response time is defined as the number of kinetic windows required to reach the pooled average basal-normalised non-mitochondrial OCR (0.287). Annotations represent the results of post-hoc TukeyHSD tests relative to 6 mM-treated samples: **, P < 0.01; *, P < 0.05; ns, not significant (P > 0.05). N = 5-7 wells.

Like our FCCP data, our results are also qualitatively comparable to those reported in other studies. With the optimal concentration of sodium azide (24 mM), OCRs were reduced to 24.2 % of MQ water controls, compared to approximately 28.5 % in Luz et al.^16^, approximately 14.3 % in Koopman et al.^21^ and approximately 20 % in Ng and Gruber^22^, indicating that our assay is able to accurately capture non-mitochondrial oxygen consumption in *C. elegans*.

### The effect of FCCP on sodium azide-treated OCRs

Respirometric experiments in intact cells are typically performed with the sequential addition of respiratory inhibitors^13^, as this allows multiple parameters of respiratory function, such as ATP synthase-linked respiration, maximal respiration and the reserve capacity, and the rate of non-mitochondrial respiration, to be assessed on a per-sample basis. However, this may not be possible in *C. elegans*, as early *C. elegans* Seahorse respirometry studies reported that the sequential use of FCCP and sodium azide increased the non-mitochondrial OCR relative to estimates derived using sodium azide alone^19^, though a later attempt to replicate this result found no such difference^21^. Given this ambiguity, we sought to determine whether non-mitochondrial OCRs were affected by FCCP supplementation using our assay. Therfore, we tracked OCRs in *C. elegans* samples treated with 24 mM sodium azide following prior treatment with 25 µM FCCP, prior treatment with 0.2 % v/v DMSO (solvent controls), or no prior treatment (untreated controls).

Our results indicated that sodium azide-treated OCRs differed depending on prior treatment (F_2, 44_ = 4.18, P = 0.0218), with OCRs significantly elevated in the FCCP-treatment group relative to the untreated group (Figure 6A, B). In contrast, DMSO-treated samples displayed OCRs intermediate to the untreated and FCCP-treated groups (both P > 0.234 relative to DMSO controls; TukeyHSD), suggesting that sodium azide-treated OCRs were elevated by the combined effects of DMSO and FCCP. It is unclear whether previous attempts to determine the impact of FCCP on sodium azide-treated OCRs controlled for DMSO^19,21^, which may be the cause of their discrepancy. Regardless of the possible role of DMSO, however, our results suggest that examining non-mitochondrial OCRs with sodium azide immediately following FCCP-induced maximal respiration will lead to an over-estimation of non-mitochondrial oxygen consumption and thus should be avoided when performing respirometry in *C. elegans*.

**Figure 6.**
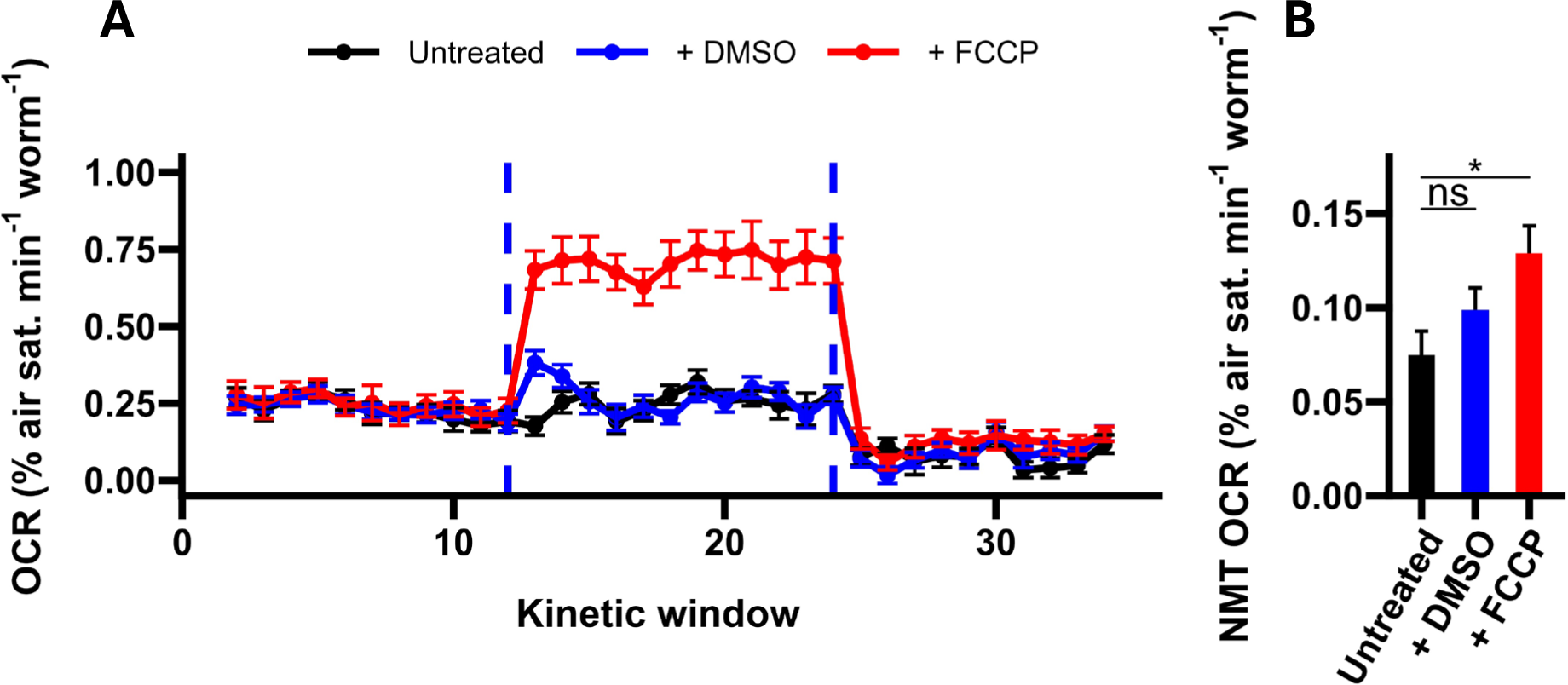
The impact of FCCP on non-mitochondrial OCRs. **A,** Oxygen consumption rate (OCR) traces of worms treated with sodium azide following prior treatment with 25 µM FCCP, prior treatment with 0.2 % v/v DMSO, or no prior treatment. FCCP and sodium azide treatments are denoted by the dashed lines. **B,** Non-mitochondrial (NMT) OCRs of untreated, DMSO-treated and FCCP-treated samples. NMT OCRs were calculated as the average OCR following the injection of sodium azide (kinetic windows 13-24). Annotations represent the results of post-hoc TukeyHSD tests relative to untreated controls: *, P < 0.05; ns, not significant (P > 0.05). N = 15-17 wells.

### General comparisons with standard methods of *C. elegans* respirometry

The assay described here allows for cost-effective, repeated OCR measurements of *C. elegans* samples with a moderate throughput. In general outline this assay resembles Seahorse XF analyser-based respirometry; oxygen levels are measured using phosphorescent oxygen sensors in a 96-well plate format, with OCRs quantified over multiple periods of oxygen consumption (measuring) and re-oxygenation (shaking). However, the major strength of this assay relative to typical high-throughput microplate-based respirometry systems is that it is comparatively inexpensive, particularly on a per-assay basis^33^.

While we have validated our assay for qualitative accuracy in day 1 adults, we emphasise that this technique is not appropriate for obtaining quantitatively accurate measurements of the *C. elegans* OCR. Our experiments were conducted without the use of a mineral oil or adhesive covering, as it was necessary to keep the wells open for re-oxygenation following each measurement period. As such, OCRs calculated using this assay are offset by oxygen diffusion from the environment, which worsens during periods of high oxygen consumption (Figure S3A). Seahorse XF respirometers are also open systems and experience oxygen diffusion^13^, but obtain quantitatively accurate OCRs by correcting raw [O_2_] data for oxygen ingress using an algorithm which employs calibration data from Clark-type respirometers^49^. We did not develop any method for the post-hoc correction of our [O_2_] data, and instead minimised the issue of oxygen diffusion by calculating OCRs over a relatively short period (60 s) in comparison to the standard 2-3 min period used by Seahorse XF analysers^13^ (Figure S3B-C). This improved the goodness-of-fit of linear regressions during periods of high (maximal) oxygen consumption relative to those obtained over longer periods (Figure S3D), but did not eliminate differences in goodness-of-fit between high (maximal) and low (basal and non-mitochondrial) oxygen consumption. As such, this technique should be used for comparative purposes only.

Since our sequential drug treatment experiments suggest that FCCP cannot be used in conjunction with sodium azide, we anticipate that others wishing to make use of this assay will likely follow the same structure used in our titration experiments, with a total assay length of approximately 82 minutes. Sample preparation and other pre-assay steps, which took no more than one hour, brought the total length of these experiments to approximately 2-2.5 h, comparable to the length of published *C. elegans* microplate-based respirometry protocols^20–24^, and easily suitable for multiple experiments in a single work day.

While maintaining a constant number of measurements (six) and measurement interval (20 s), we were able to measure a total of 10 samples per assay using our basic OCR protocol and script. This throughput is similar to the 8-well Seahorse XFp^23^, and significantly higher than the two-chamber Oxygraph-2k^33^, but lower than the 24-well Seahorse XF24^19,20^ and 96-well XF96^21^. However, when considering overall throughput, it is notable that pooling data from replicate Seahorse experiments in *C. elegans* is considered inappropriate in published protocols^21^. This largely results from the fact that Seahorse systems were developed for cell cultures and possess heating elements fixed at 37 °C^21^. As this temperature presents an acute heat stress for *C. elegans*^50^, this heating element must be turned off prior to use, allowing the internal temperature of the system to equilibrate to the laboratory temperature (20 °C) but leaving it free to increase by as much as 5 °C during operation^20–24^. This limitation did not apply here as we opted to use the BMG FLUOstar’s modifiable heating element to maintain an internal temperature of 25 °C (its lowest setting), which typically deviated by < 1 °C during an experiment. While it could be argued that 25 °C is a sub-optimal temperature for *C. elegans* respirometry, as incubation at this temperature is associated with alterations in mitochondrial dynamics and elevated metabolic rates^51,52^, these phenotypes are associated with long-term incubation at this temperature. We argue that if short-term alterations in temperature are predicted to significantly alter the *C. elegans* OCR, then it is better to maintain a constant elevated temperature than to allow temperatures to slowly increase over the course of an experiment, possibly leading to erroneous estimations of respiratory parameters calculated from periods of oxygen consumption under different ambient temperatures (e.g. the reserve capacity: maximal minus basal OCR). Performing our experiments at a constant incubation temperature, combined with the inherent reduction of well-well and run-run variability related to the internal referencing of the OxoPlate sensors, enabled us to pool the data from multiple experiments for analysis. When taking this into account, the practical throughput of this assay is more similar to the Seahorse XF24 and XF96 analysers, making it easily suitable for larger experiments with multiple conditions.

### Conclusions

Here we describe the development and validation of a cost-effective assay for *C. elegans* respirometry which does not require access to specialised respirometry equipment. We validate our assay for comparative accuracy in day 1 adults by demonstrating the linearity of basal OCRs in samples with highly variable numbers of animals, and by examining the effects of the respiratory inhibitors FCCP and sodium azide in titration experiments. We then apply our assay to demonstrate that prior treatment with FCCP elevates sodium azide-treated OCRs, supporting previous *C. elegans* respirometry data, and strongly suggesting that estimates of the maximal and non-mitochondrial OCR must be obtained separately when working with *C. elegans*.

While our experiments were limited to wild-type day 1 adults, this assay could be easily adapted for use in animals of different ages or genetic backgrounds by simply replicating our sample size and titration experiments. This assay will be useful for researchers who wish to perform respirometry in *C. elegans* in a cost-and time-efficient manner.

## Acknowledgements

We acknowledge support from the Biotechnology and Biological Sciences Research Council (BBSRC) through grant BBSRC(BB/V011243/1), and Dr John Stolz through a PhD studentship awarded to Nathan Dennis. Some strains were provided by the CGC, which is funded by NIH Office of Research Infrastructure Programs (P40 OD010440).

## Supplementary Information

### OCR protocol script

All OCR assays were conducted using the time-resolved fluorescence protocol described in Table 2 of the main text with the following BMG FLUOstar script:

### BMG FLUOstar OCR script

st1: = “OxoPlate protocol” #Pre-configured OxoPlate protocol

NumberOfReadings: = n

wait for 10 min

for i=1 to NumberOfReadings do begin

ID1:=”Reading “ i

R_run “<ST1>”

If i = 3 Then begin

Ask “Add drug A and click Yes to continue “ (”Continue”)

End

If i = 6 Then begin

Ask “Add drug B and click Yes to continue “ (”Continue”)

End

Next i

Pauses were implemented every 3 cycles to allow for the plate to be ejected from the reader and treated with respiratory inhibitors. The initial wait period (10 minutes) was included to ensure the temperature of the wells had equilibrated with the internal temperature of the plate reader, and was estimated by examining the stabilisation of oxygen levels in solutions of Milli-Q water following a shift from 20 °C (laboratory temperature) to 25 °C (the internal temperature of the plate reader; Figure S4), which occurred within approximately 10 minutes of measurement. For each assay, the NumberOfReadings (n) was changed as necessary. Sample size analysis was performed with n = 2; dose-response analysis with n = 6, and sequential drug addition experiments with n = 9.

### Automated OCR calculation script

To expedite the calculation of OCRs from internally referenced (I_R_) sensor responses, we created a set of functions in R v. 4.3.3 to automatically calculate oxygen concentrations, fit linear regressions within user-defined measurement windows, extract the regression coefficient, multiply the result by −1, and normalise to worm number (or any alternative normalisation parameter). This script requires that the data be formatted with a single “time” column (which must be in lowercase and formatted in seconds), and multiple I_R_ data columns with the name of the condition specified first, followed by an underscore and a designation for the replicate (see Figure S5 for an example data sheet). This script also requires pre-determined calibration constants derived from oxygen-free (k_0_) and oxygen-saturated (k_100_) calibration solutions (see OxoPlate sensor calibration), and the pre-installation of the *tidyverse* and *forstringr* packages. For the OCR calculation script, the OCR calculation window is set by the interval value (the number of measurements for regression-fitting), and the total length of each kinetic window (as set by the number of cycles in the BMG FLUOstar protocol) is set by the step value.

### [O_2_] calculation function

k0 <-#User-defined k_0_ value

k100 <-#User-defined k_100_ value

o2.fun <-function(data, k0, k100){

df <-cbind(data[,1], 100*(k0/data[,-1]-1)/(k0/k100-1))

return(df)

}

### Regression-fitting and differentiation function

get_slope <-function(x, y) {

lm_model <-lm(y ∼ x) #Fits linear regression

return(coef(lm_model)[2]) #Extracts regression coefficient

}

### OCR function

OCR.fun <-function(data, interval, step, worm.no) {

slopes <-numeric() #Initialises empty vector to store OCRs for each kinetic window

col_names <-character() #Initialises empty vector to store sample name information

interval_numbers <-numeric() #Initialises empty vector to record the kinetic window

interval_number <-1 #Sets kinetic window number to 1 prior to running the loops

for (i in seq(1, nrow(data), by = step)) { #Loop to subset the data for each kinetic window

subset_data <-as.data.frame(data[i:(i + interval - 1),])

time_subset <-subset_data$time #Extracts time column for calculating OCRs

for (col in 2:ncol(subset_data)) { #Loop to fit regressions to each sample within each window

slope <-get_slope(time_subset, subset_data[, col])

slopes <-c(slopes, slope / worm.no[col - 1])

col_names <-c(col_names, names(subset_data)[col])

interval_numbers <-c(interval_numbers, interval_number) #Records the kinetic window

}

interval_number <-interval_number + 1 #Moves to the next kinetic window

}

result <-data.frame(cycle = interval_numbers, #Combines the data into a single data frame

rep = col_names, der = -slopes * 60, #Expresses OCRs per minute

condition=str_extract_part(col_names, “_”, before = T)) #Extracts condition identifiers

return(result) #Returns the results as a data frame for analysis or export

}

## Supplementary tables

**Table S1.**
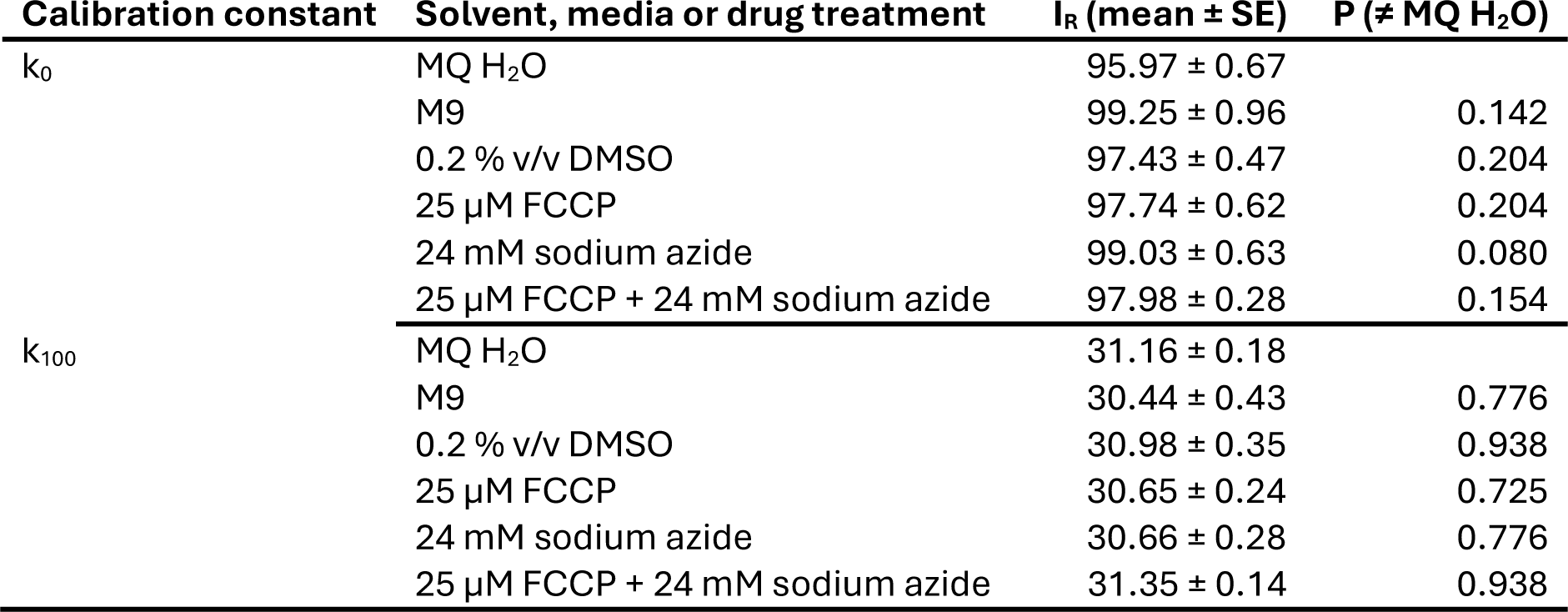
Calibration constants for all media, solvent and drug treatments. P values represent the results of FDR-corrected Student’s t tests relative to Milli-Q (MQ) H_2_O controls. N=3-4.

## Supplementary figures

**Figure S1.**
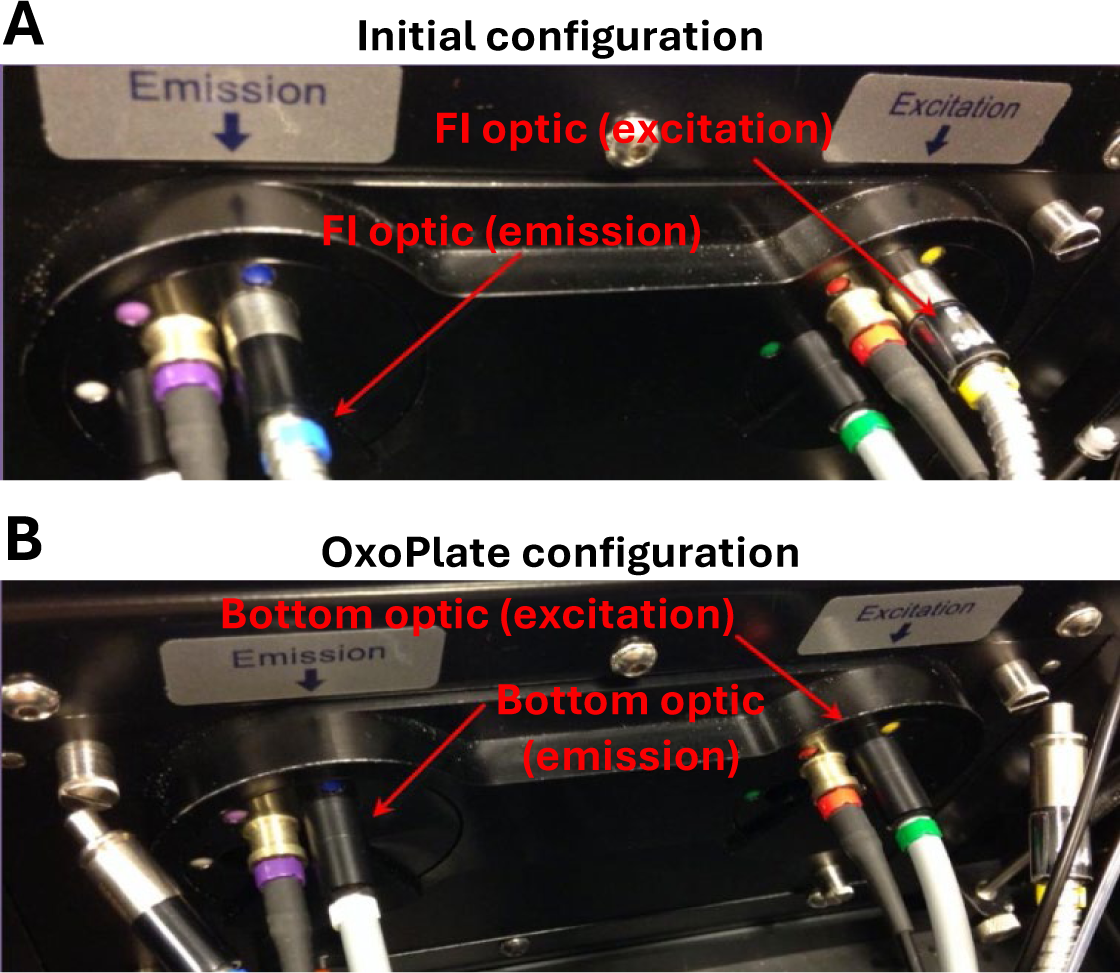
Light guide configuration for bottom-optic time-resolved fluorescence measurements using the BMG FLUOstar. **A,** Default light guide configuration with blue and yellow fluorescence intensity (FI) light guides. **B,** Corrected configuration in which the yellow and blue FI light guides have been replaced with the white and green bottom-optic light guides, leaving the FI light guides disconnected. Both Images were provided by PreSens.

**Figure S2.**
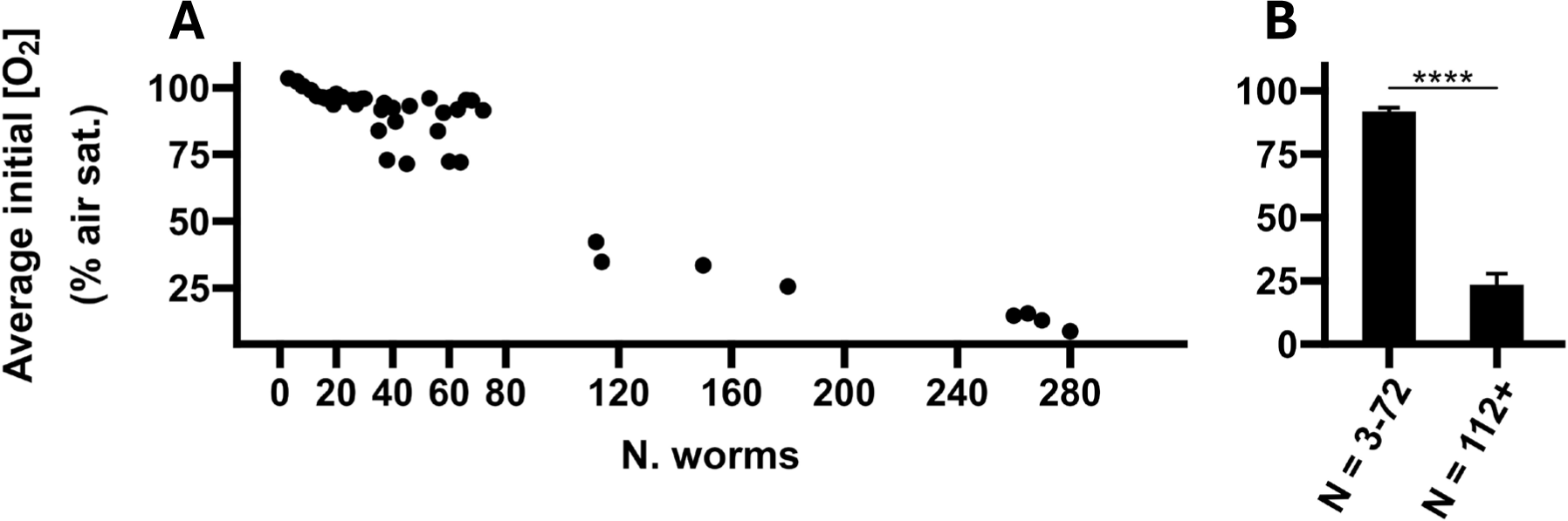
The impact of sample size on average starting [O_2_]. **A,** Average first-measurement [O_2_] in wells containing 3-280 animals. **B,** Pooled average first-measurement [O_2_] over the linear OCR range (3-72 animals per well) compared to the non-linear range (112-280 animals per well). N = 40 wells. Annotations represent the results of ANOVA. ****, P < 0.0001.

**Figure S3.**
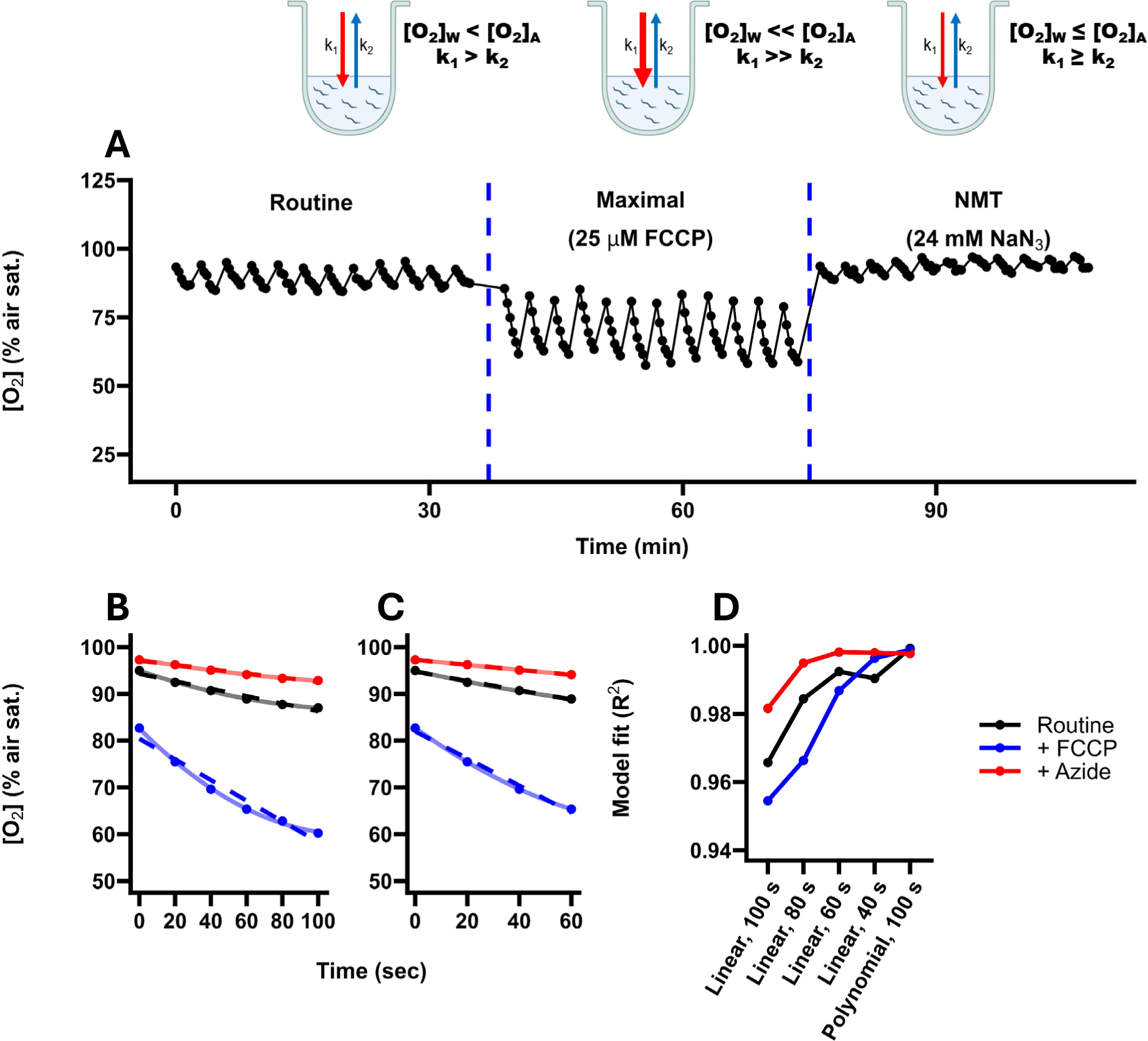
Defining optimal windows for the calculation of the *C. elegans* OCR. **A,** Simplified models of oxygen diffusion under periods of moderate (basal), high (maximal) and low (non-mitochondrial, NMT) oxygen consumption in uncovered OxoPlate wells with an accompanying average [O_2_] trace of *C. elegans* samples sequentially treated (dashed lines) with FCCP and sodium azide (NaN_3_; re-produced from Figure 6; N = 15 wells). During periods of low oxygen consumption, the oxygen concentration within each well ([O_2_]_W_) is similar to the oxygen concentration of the ambient air ([O_2_]_A_), resulting in similar rates of oxygen diffusion (k_1_ and k_2_) between the environment and the contents of each well. In contrast, during periods of high oxygen consumption, the [O_2_]_W_ drops significantly below the [O_2_]_A_, causing a large net influx of oxygen into the contents of each well, masking the biological OCR. These models are simplified to only consider the contents of each well and the ambient air. However, the material of the OxoPlates likely acts as a separate oxygen reservoir and may need to be accounted for in an accurate model of oxygen diffusion. **B-C,** Oxygen concentrations averaged over each kinetic window displayed in panel A during basal, maximal and non-mitochondrial (NMT) respiration, with fitted curves of 2^nd^-order polynomials (solid lines) and linear regressions (dashed lines). Curves are fitted to the full 100 seconds in B, and the first 60 seconds in C. **D,** Goodness-of-fit (R^2^) of linear models using subsets of the full measurement window (100 s) in 20 s increments, compared to the goodness-of-fit of 2^nd^-order polynomials fit to the full measurement window.

**Figure S4.**
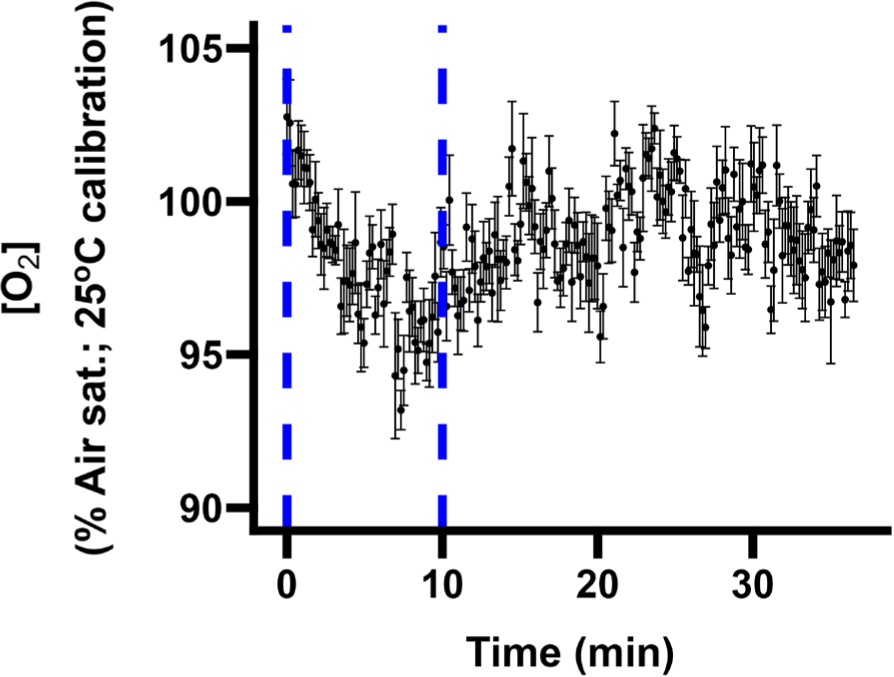
Estimate of temperature equilibration time following a shift from 20 °C to 25 °C. Temperature equilibration time was estimated based on the approximate length of time required for oxygen levels (quantified based on a two-point calibration at 25 °C) in solutions of Milli-Q to cease declining following transfer from 20 °C to 25 °C. N = 5 wells.

**Figure S5.**
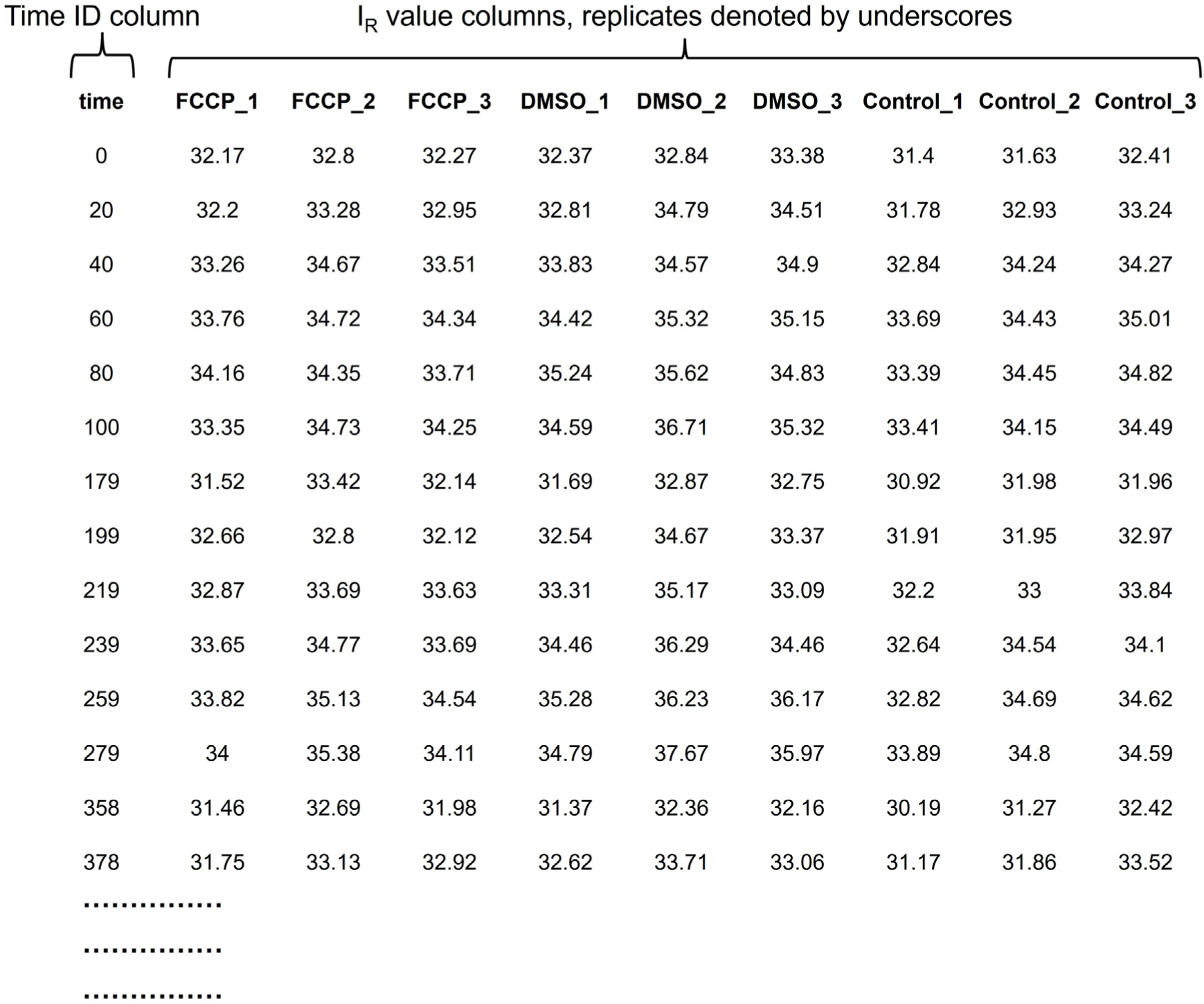
Example data for automatic OCR analysis. The time identifier must be labelled “time” (in lowercase) and formatted in seconds. Each data column must be formatted with the name of the condition first (here FCCP, DMSO or Control) followed by a designation for the replicate (numbers or letters) separated by an underscore, i.e. FCCP_1 refers to the first replicate of the condition “FCCP”.

